# Collective microfibril sliding underlies plant cell wall creep

**DOI:** 10.64898/2026.03.24.713984

**Authors:** Changhao Li, Jingyi Yu, K. Jimmy Hsia, Daniel J. Cosgrove, Sulin Zhang

## Abstract

Plant cell growth depends on slow, irreversible creep of the fibrous cell wall stretched by turgor, yet the mechanics of creep—and how it differs from elasto-plastic deformation—remains uncertain^1–3^. Using multiscale modeling, we show how wall creep emerges from elementary sliding of cellulose microfibrils along their contact interfaces. A dominant sliding mode involves localized dislocation-like defects that stochastically nucleate at laterally bonded microfibril contacts and then glide along the interface by surmounting discrete energy barriers. Embedding these sliding events into Monte Carlo simulations of stretched cellulose networks recapitulates whole-wall creep kinetics. Elevated stress lowers sliding energy barriers and triggers rapid irreversible deformation characteristic of plastic flow. Compared with elasto-plastic stretching, creep induces less microfibril reorientation but promotes bundling, thereby reducing subsequent yielding. These results clarify the microfibril rearrangements underlying cell wall creep and show how they redistribute stress to permit sustained wall expansion at steady turgor without compromising structural integrity.

## Main Text

Plant cells commonly enlarge more than 100-fold in volume during growth, generating diverse shapes at the cellular, organ, and whole-plant scales^2,3^. This process, which underlies both the impressive height of redwood trees and the rapid growth of weeds, entails sustained irreversible cell wall enlargement by turgor-driven creep of the growing cell wall, a dynamic nanostructured polymeric material that uniquely combines strength, extensibility, and mechanical integrity^1,4^ in ways seldom realized in engineering materials^5,6^. A century of materials research has elucidated multiscale mechanisms of creep in engineering materials^7^, whereas the mechanics of cell wall creep, essential to plant growth, is largely unknown, despite extensive biomechanical modeling of plant growth and cell walls^8–12^.

At the molecular level, the growing wall consists of a strong noncovalent network of crystalline cellulose microfibrils (CMFs) embedded in a hydrated pectin–hemicellulose matrix^1,13^, as shown in AFM images (**Fig. 1a**). At short timescales, the wall responds to tensile loading through a combination of reversible (elastic) and irreversible (plastic) deformation^14^. Such elasto-plastic responses underpin some recent models of plant mechanobiology and morphogenesis^10,15^, yet growth more closely resembles slow wall creep, which generates most of the surface expansion during growth and is mechanically driven by steady wall stresses generated by turgor pressure^3,16^. A recent coarse-grained molecular dynamics (CGMD) model^8,17^ clarified how CMFs can assemble into a strong yet extensible load-bearing network in which tensile forces are transmitted predominantly through noncovalent CMF-CMF contacts, whereas matrix polysaccharides (xyloglucan, pectin) modulate CMF spacing and network formation, but are predicted to carry comparatively little tensile load. The model, which is based on the cross-lamellate structure of epidermal walls, attributes wall elasticity and plasticity largely to microfibril bending and sliding, respectively, within the cellulose network, but does not account for the long-timescale process of creep.

**Fig. 1.**
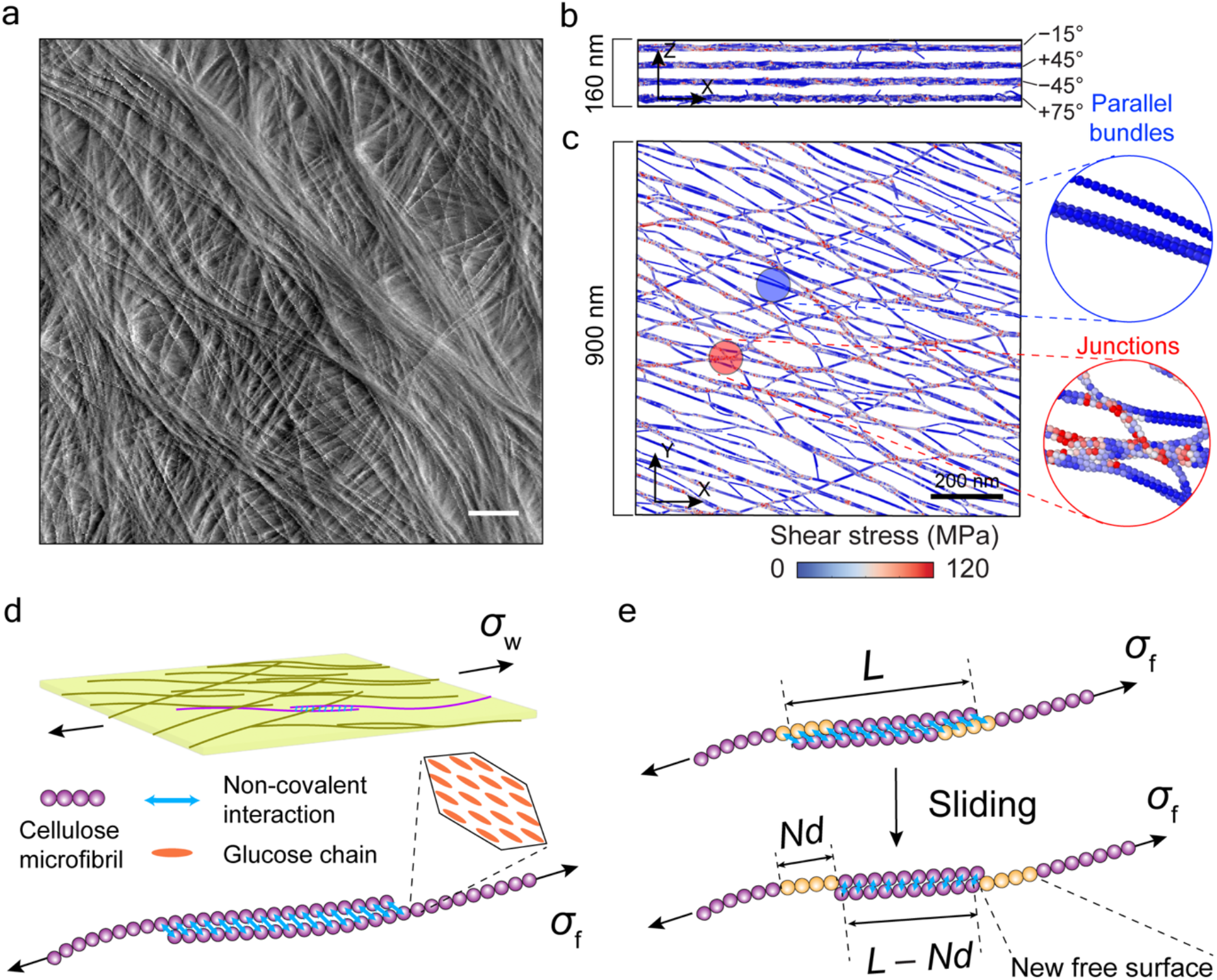
Force transmission and CMF sliding in the hierarchical structure of multi-lamellate cell walls. **(a)** An AFM Peakforce error image of an onion outer epidermal wall, showing multiple lamellae with anisotropic CMF organization. **(b)** Side view of the equilibrated coarse-grained configurations for a four-lamella wall model under 5% tensile strain, composed of lamellae with CMF directions −15°, +45°, −45°, +75° relative to the x-axis (stretch direction). **(c)** Top view of equilibrated configurations for the −15° lamella under 5% tensile strain, colored by per-bead interfacial shear stress magnitude. Shear stresses are concentrated in irregularly packed regions such as junctions and free ends, whereas the bulk regions of regular CMF bundles exhibit minimal shear stress. **(d-e)** Schematics of force transmission and inter-microfibril sliding during wall deformation. In (d), interconnected CMF networks act as the major load-bearing structures; in (e), when the fibril stress exceeds the creep threshold, a new free surface is created by a sequence of elementary sliding events between CMFs. Initial equilibrium state with contact length *L* (top), and final equilibrium state after sliding with contact length *L* − *Nd* (bottom) are schematically sketched, where *Nd* is the cumulative sliding distance after *N* elementary sliding events. Scale bars, 200 nm.

In growing plant cells, wall creep is mechanically powered by turgor pressure and biochemically modulated by the wall-loosening activity of α-expansin proteins acting downstream of the hormone auxin and other developmental regulators^18–20^. Here we model the *mechanical* process of creep, i.e., stress-driven wall yielding in the absence of *biochemical* loosening agents such as α-expansin, to establish a baseline molecular-mechanical mechanism that later can be augmented with expansin action. We extend the CGMD framework to reach much longer timescales^21–23^ by incorporating CMF sliding kinetics based on computed sliding energy barriers. Our extended simulations show how collective CMF sliding results in wall creep, simultaneously remodeling the cellulose network and redistributing stresses while preserving the wall’s load-bearing architecture. We then contrast the structural effects of creep with those of elasto-plastic stretching, documenting distinctive changes in the wall.

We consider CMF sliding as the dominant rate-limiting molecular process underlying irreversible extension of epidermal walls, which limit growth of many plant organs. This concept is supported by AFM imaging visualizing CMF-CMF networks and sliding in epidermal walls^24^, coarse-grained modeling of the structure and tensile mechanics of epidermal cell walls^8,17^, experimental tests indicating limited roles of pectin and xyloglucan in tensile mechanics of epidermal walls^14,16^, and extensive evidence that cellulose is the major load-bearing component of cell walls. Cell wall stretching activates multiple elementary CMF motions, including straightening, curving, reorientation, and sliding^8,24^. Our cross-lamellate CGMD model (**Supplementary Note 1**) includes four anisotropic lamellae with initial cellulose orientations averaging −15°, +45°, −45°, +75° (**Fig. 1b**). Shear stresses arise at CMF interfaces within the lamellae (**Fig. 1c**). When the wall is stretched by wall stress *σ*_w_, tensile forces are transmitted through limited load-bearing contacts in the cellulose network, generating local fibril stresses *σ*_f_ (**Fig. 1d**) 10-to 100-fold higher than *σ*_w_. This produces interfacial shear stresses sufficient to drive CMF sliding^8^ (**Fig. 1e**), dissipating mechanical energy by creating new free surfaces. During sliding, noncovalent bonds between CMFs dynamically break and reform, releasing local stresses and transferring loads to neighboring CMFs, but transient states during sliding are unknown. We follow a bottom-up strategy to investigate how CMFs slide and how sliding orchestration results in whole-wall creep.

### Elementary sliding kinetics of laterally-bonded cellulose microfibrils: a two-chain model

We begin by modeling sliding between a pair of CMFs with contact length *L* and tensile stress *σ*_f_ imposed at their ends (**Fig. 1e**; key parameters summarized in **Table 1**). Each CMF is represented as a bead-and-spring (coarse-grained) chain that captures the essential physical properties of CMFs^8^. The sliding process can be viewed as a sequence of discrete thermally-activated transitions between local equilibrium states^25^, driven by mechanical stresses^26^ and kinetically limited by energy barriers between states. This is analogous to biochemical reactions limited by activation energy barriers. Each elementary sliding event corresponds to the chains shifting in register by one bond length *d* (equivalent to bead diameter) under constant stress *σ*_f_. The thermodynamic driving force for sliding arises from the energy difference between the two states (**Fig. 1e**), which depends on fibril stress *σ*_f_. The mechanical work gained by advancing one registry length scales with applied fibril force times displacement, whereas the interfacial cost scales with newly created free surface area times surface energy density. At relatively low stresses, the work done per sliding event, estimated as *σ*_f_*d*^3^, is insufficient to compensate the energy cost to overcome non-covalent interactions *γd*^2^ (*γ* is the surface energy density, which depends on interaction forces between cellulose beads^8^) and the increase in strain energy. Under this condition, forward sliding is thermodynamically unfavorable. Neglecting secondary strain energy contributions, *σ*_c_ ~ *γ*/*d* defines the stress threshold where sliding becomes thermodynamically favorable, marking the onset of stress-driven microfibril sliding.

**Table 1.** Definition and range of important quantities.

Macroscopic quantities (tissue or cellular level)
| Symbol | Definition | Range | Unit |
| --- | --- | --- | --- |
| $\varepsilon_w$ | Normal strain of cell walls in the stretching direction | 0–50% | 1 |
| $\sigma_w$ | Normal stress applied to cell walls in the stretching direction | 0–10 | MPa |
| $Y$ | Wall-scale creep threshold | $\sim 0.2$ | MPa |
| | Wall-scale plastic yield stress | $\sim 4$ | MPa |
| $\dot{\varepsilon}_w$ | Wall-scale creep strain rate | $10^{-5}$–$10^{-2}$ | $\text{min}^{-1}$ |

Microscopic quantities (molecular level)
| Symbol | Definition | Range | Unit |
| --- | --- | --- | --- |
| $\sigma_f$ | Normal stress applied to CMFs in the chain direction | 0–1000 | MPa |
| $\tau$ | Shear stress on a coarse-grained CMF bead | 0–200 | MPa |
| $d$ | CMF diameter (bond length) | 3 | nm |
| $\gamma$ | Surface energy of CMFs | $\sim 0.10$ | N/m |
| $\sigma_c$ | CMF-scale creep threshold | $\sim 30$ | MPa |
| $\sigma_p$ | CMF-scale plastic yield threshold | $\sim 300$ | MPa |
| $\Delta E$ | Energy barrier of CMF sliding | 0–200 | $k_B T$ |
| $k_B T$ | Energy of thermal fluctuations (under 300 K) | $\sim 0.025$ | eV |
| $\dot{\varepsilon}_f$ | Sliding strain rate for a CMF pair | $10^{-5}$–$10^{-2}$ | $\text{min}^{-1}$ |
| $L$ | Contact length between a pair of CMFs | 0–850 | nm |

Above *σ*_c_, sliding is energetically favored but kinetically hindered by sliding barrier Δ*E*, largely stemming from adhesion of CMFs. We used the nudged elastic band (NEB) method^27^ to compute the minimum energy path and the energy barrier Δ*E* for sliding at prescribed fibril stresses *σ*_f_ and contact lengths *L*. Two distinct sliding mechanisms emerge: For small *L*, all beads within the contact region shift coherently by one bead diameter at a time (*uniform sliding*, **Fig. 2a, Video S1**). As *L* increases, the sliding barrier increases, uniform sliding becomes kinetically disfavored, and a second mechanism, which we call *dislocation-mediated sliding*, begins to dominate (**Fig. 2b, Video S2**). Localized dislocation-like defects in CMF bonding nucleate at microfibril ends where shear stresses concentrate due to shear-lag effects^28^, resulting in stochastic, thermally-generated compressive defects which then glide down the length of the microfibril. Short contacts favor uniform sliding whereas long contacts favor dislocation-mediated sliding because Δ*E* scales differently with *L* for the two mechanisms and the energy barrier for dislocation sliding plateaus at L equivalent to the dislocation length (**Extended Data Fig. 1**). In effect, the dislocation mechanism provides a lower energy pathway for sliding.

**Fig. 2.**
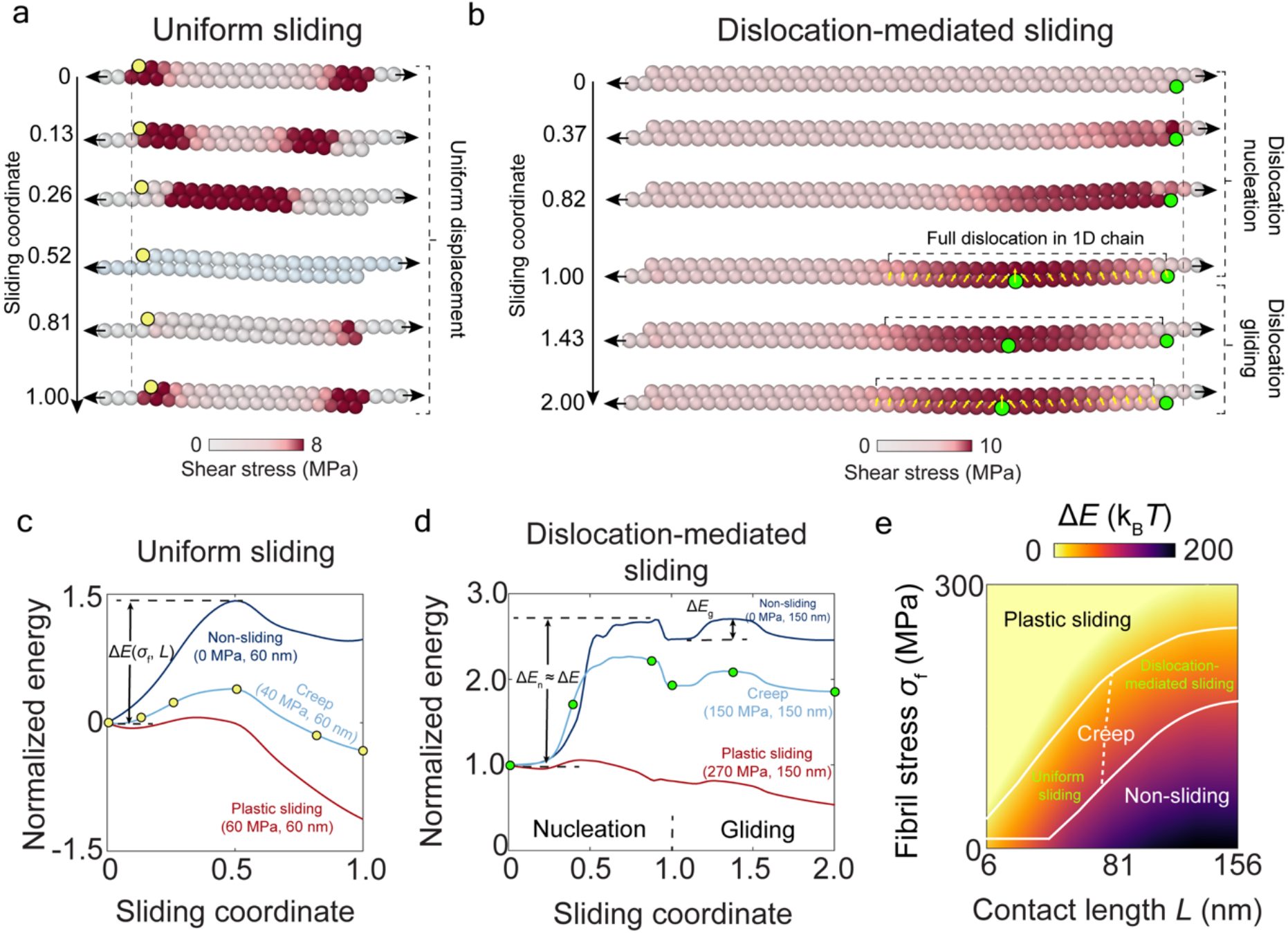
Sliding kinetics of CMF pairs depends on contact length and fibril stress. **(a-b)** Representative intermediate states for uniform sliding (a, *L* = 60 nm, *σ*_f_ = 40 MPa) and dislocation-mediated sliding (b, *L* = 150 nm, *σ*_f_ = 150 MPa). In (a), left free ends are colored by yellow dots to highlight the progress of uniform sliding. In (b), dislocation patterns are labeled by yellow solid arrows. One free end and the center of the dislocation are highlighted by green dots to visualize the dislocation nucleation and gliding process. See Methods for the definition of sliding coordinates. Note that NEB methods can capture intermediate states even if sliding is not thermodynamically favorable. **(c-d)** Minimum energy paths for uniform sliding (c, *L* = 60 nm, *σ*_f_ = 0, 40, 60 MPa) and dislocation-mediated sliding (d, *L* = 150 nm, *σ*_f_ = 0, 150, 270 MPa). Yellow dots in (c) and green dots in (d) label corresponding transition states in (a) and (b) respectively, with different sliding coordinates. In (d), dislocation nucleation and gliding overcome separate energy barriers Δ*E*_n_ and Δ*E*_g_, respectively. Energies are normalized by 2*D*_01_ = 90 *k*_B_*T*, where *D*_01_ is the non-bonded interaction strength between two beads. **(e)** Phase diagram delineating regions of non-sliding (elastic deformation only), creep, and plasticity, separated by white solid lines. The creep region is further separated by two distinct sliding mechanisms: uniform sliding at low contact lengths and dislocation-mediated sliding at high contact lengths.

Different from the effect of contact length *L*, changing fibril stresses *σ*_f_ does not alter the sliding mechanism but tilts the energy landscape to influence sliding kinetics. Elevated stresses decrease the sliding barrier for both sliding mechanisms, but their energy landscapes differ (**Figs. 2c, 2d**). For dislocation-mediated sliding, system energy must exceed the nucleation barrier Δ*E*_n_ to generate the dislocation which glides earthworm-like into the overlapping region (**Extended Data Fig. 2**), overcoming successive gliding barriers Δ*E*_g_ (**Fig. 2d**); because Δ*E*_n_ is substantially larger than Δ*E*_g_, dislocation nucleation is the rate-limiting step. The stress at which Δ*E* vanishes defines a plastic yield threshold *σ*_p_, above which CMF sliding is rapid and spontaneous. Notably, the plastic threshold *σ*_p_ is much larger than the creep threshold *σ*_c_ (**Table 1**), distinguishing the slow stochastic creep regime (*σ*_c_ < *σ*_f_ < *σ*_p_) from the rapid plastic sliding regime *σ*_f_ ≥ *σ*_p_ with distinct time scales of wall expansion. This is illustrated in a phase diagram showing three regimes of CMF motions (**Fig. 2e, Extended Data Fig. 3**): i) a non-sliding regime (*σ*_f_ < *σ*_c_), where CMFs undergo purely elastic deformation without sliding (neglecting effects of CMF network topology and matrix viscosity); ii) an intermediate creep regime (*σ*_c_ < *σ*_f_ < *σ*_p_) including uniform sliding at short contact lengths and dislocation-mediated sliding at longer lengths, and iii) a plastic regime (*σ*_f_ > *σ*_p_) where rapid plastic yielding dominates as the barrier Δ*E* is reduced.

This minimal two-chain model captures the essential features of creep even without considering complex CMF networks. The model also sheds light on “stick-slip” dynamics of wood plasticity^29^ where repetitive accumulation and release of interfacial stresses accompany irreversible stretches and where transient states at the onset of stress release are poorly understood due to timescale limitations of experiments and conventional MD simulations. Our model fills this gap by capturing the complete dynamics of stick-slip motions, commonly observed in cellulose-based materials^29–31^.

### Collective microfibril sliding as the basis of wall creep

To connect the sliding of CMF pairs to creep of whole walls, we employed the Monte Carlo (MC) method^32^ where each MC step corresponds to a single sliding event between CMF chains (**Supplementary Note 2**). The MC sampling rule was based on Arrhenius-type kinetics for thermally-activated barrier-crossing processes: sliding rate 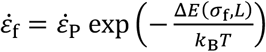, where 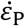 is the maximum sliding strain rate in the plastic regime, representing the upper limit of 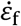, and the energy barrier Δ*E*(*σ*_f_, *L*) is interpolated from the NEB calculations in **Fig. 2e**. After each MC step, the system is equilibrated by CGMD before proceeding to the next MC step. This iterative sequence simulates the time course of wall creep, capturing long trajectories of thermally-activated rare sliding events with efficiency and stability. The coupled algorithm exceeds the typical sub-microsecond limit of MD simulations^33^ and can simulate creep and stress relaxation lasting more than an hour.

To parameterize this model we performed creep experiments with onion epidermal wall strips^34^ (**Fig. 3a**) to estimate 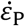 from the strain-time data at three loads (Methods). In both experiments and simulations, cell walls were stretched under constant force boundary conditions, where creep rates increase with applied stress but decay with time (**Fig. 3b**). Creep simulations capture observable collective CMF sliding and consequential structural evolution (**Video S3**). The sliding rate exhibits strong anisotropy among the lamellae and fluctuates depending on local stress distributions and microstructures (**Fig. 3c, Extended Data Fig. 4**). During creep, the −15° lamella (aligned most closely to the stretching direction) sustained the highest tensile stresses^8^ and so experienced the highest sliding rates among the four lamellae. In the −15° lamella, sliding rates gradually decayed and stabilized within 40 min (**Fig. 3d**), consistent with whole-wall responses (**Fig. 3b**). During wall creep, CMFs changed their orientation (by angle change Δ*α*) and connectivity *C* (average number of connected neighbors for each microfibril) with similar decay kinetics (**Fig. 3e, 3f**). These molecular rearrangements may gradually suppress further sliding by redistributing stresses (**Fig. 3g**) and changing local contact geometries (**Extended Data Fig. 5**).

**Fig. 3.**
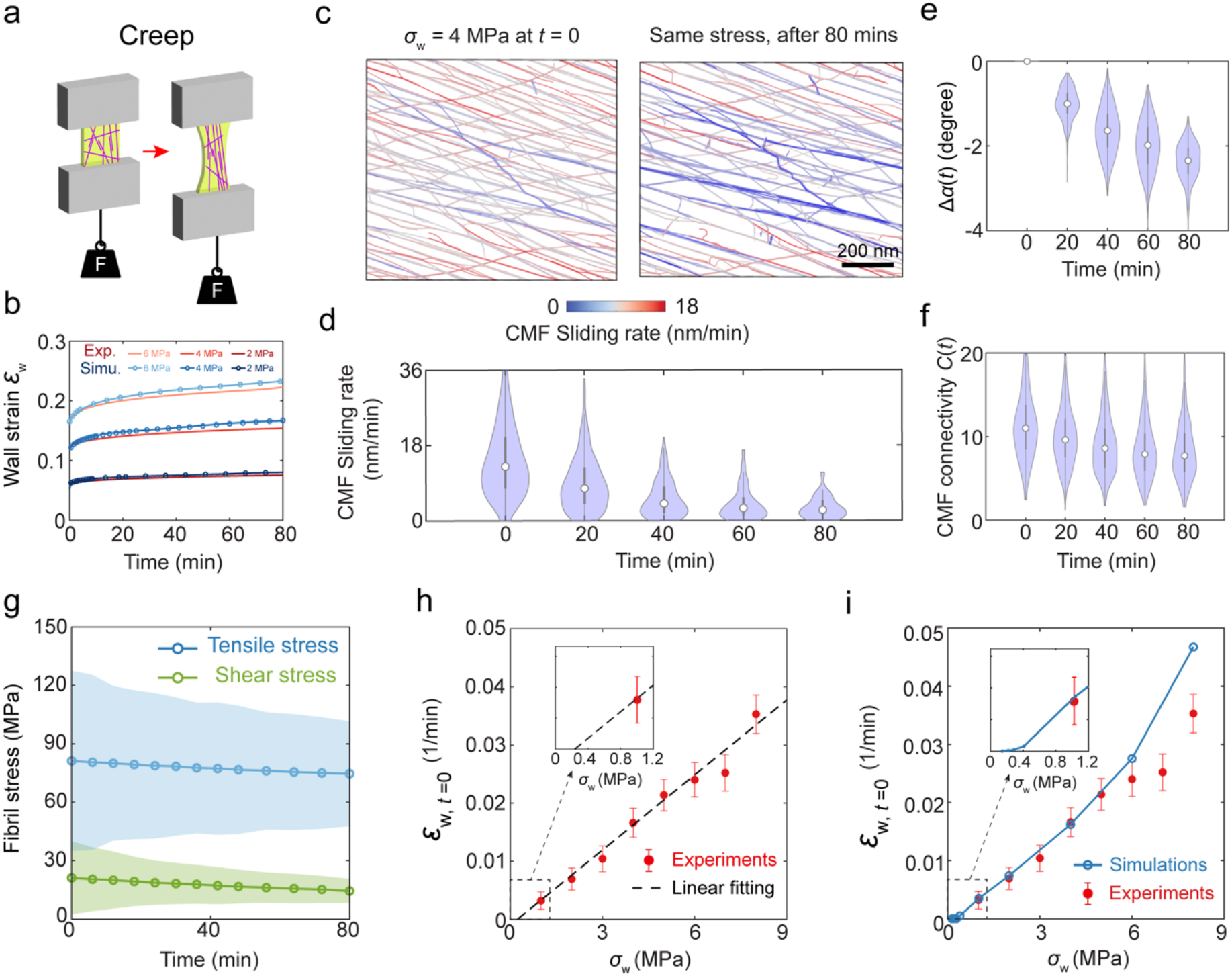
Stress and microstructure evolution during cell wall creep. **(a)** Experimental and simulation protocols of creep with constant uniaxial loading. **(b)** Time courses of wall strain under constant tensile stress *σ*_w_ = 2 MPa, 4 MPa, 6 MPa, from experimental and numerical creep tests. **(c)** Spatiotemporal evolution of CMF sliding rates 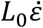 in the –15° layer for creep (*σ*_w_ = 4 MPa). Scale bar: 200 nm. **(d-f)** Violin plots showing the time evolution of sliding rate (d), reorientation angle (e), and CMF connectivity (f) for creep (*σ*_w_ = 4 MPa). **(g)** Time evolution of average CMF tensile stress and shear stress in the −15° lamella during creep (*σ*_w_ = 4 MPa). **(h-i)** At *t* = 0, experimental (h) and simulated (i) cell wall creep rates under different tensile stresses. In (h), experimental creep rates are linearly extrapolated to estimate the creep threshold. Inset: magnified view showing the creep threshold. Experimental values are means ± standard deviation of 3 measurements.

At *t* = 0, the measured and simulated creep rates increase with wall stresses, and their linear extrapolation reveals a wall-scale creep threshold *Y* ~ 0.25 MPa (**Fig. 3h, 3i**), where the creep rate nearly vanishes. This observation is consistent with experimental results^35^ and Lockhart’s proposal^36,37^ that growth rate scales with turgor pressure above a threshold. Wall stresses below *Y* generate negligible creep because most CMF pairs sustain stresses smaller than the sliding threshold *σ*_c_. The creep threshold is well below the experimentally estimated plastic yield threshold ~4 MPa^14^, confirming that creep occurs at moderate wall stresses. The physical creep rate also decays with time, coordinately with changes in CMF orientation Δα, connectivity *C*, and possibly other network properties. The two scaling relations with stress and time yield a creep law 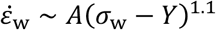, where *A*(*t*) is a time-dependent parameter reflecting the structural changes of the CMF network, and the exponent 1.1 is fitted from simulations, consistent with measured exponents of 1.0–1.5 from growth^35^ and our creep experiments (**Extended Data Fig. 6**). This creep law resembles initial and intermediate states of polymer creep^38^, where long-chain molecules undergo decayed internal relaxation driven by sustained applied tension. The network-level simulations show that CMF sliding at a broad range of rates contributes to network creep and highlights the microstructural origin of the wall-scale creep threshold *Y*, which originates from CMF-scale creep thresholds *σ*_c_. Time-dependent sliding and stress redistribution enable irreversible extension at wall stresses well below the plastic yield threshold, mimicking wall extension during turgor-driven growth^36^. In vivo, a high creep rate is sustained by the continuous action of α-expansin^18^. Our results naturally lead to a conjecture that α-expansins catalyze creep by binding CMF networks and facilitating dislocation-mediated sliding kinetics.

### Sliding-induced cellulose network rearrangement strengthens cell walls

Both creep and plasticity lead to irreversible elongation, but they differ in timescale and microstructural changes during deformation. These differences in turn lead to diverging wall structures and mechanical properties, which we assessed in the following experiments. Two identical digital walls were brought to 15% strain by either rapid elasto-plastic stretching or slow creep, followed by unloading to zero stress (**Fig. 4a**), then rapidly stretched in a second loading cycle to 20% strain (**Fig. 4b**). Creep resulted in comparatively more irreversible deformation in the first cycle, but less energy dissipation in the second cycle (**Fig. 4c** and **Video S4**). In effect, the wall remembered its past deformation, a memory encoded in its molecular structure. Slow creep leads to a distinct wall state, characterized by increased stiffness upon reloading and greater resistance to further plastic flow. This suggests that creep deformation during growth may remodel the cellulose network in a way that stabilizes its mechanics, enabling cells to maintain structural integrity while continuing to grow under persistent turgor-driven stress.

**Fig. 4.**
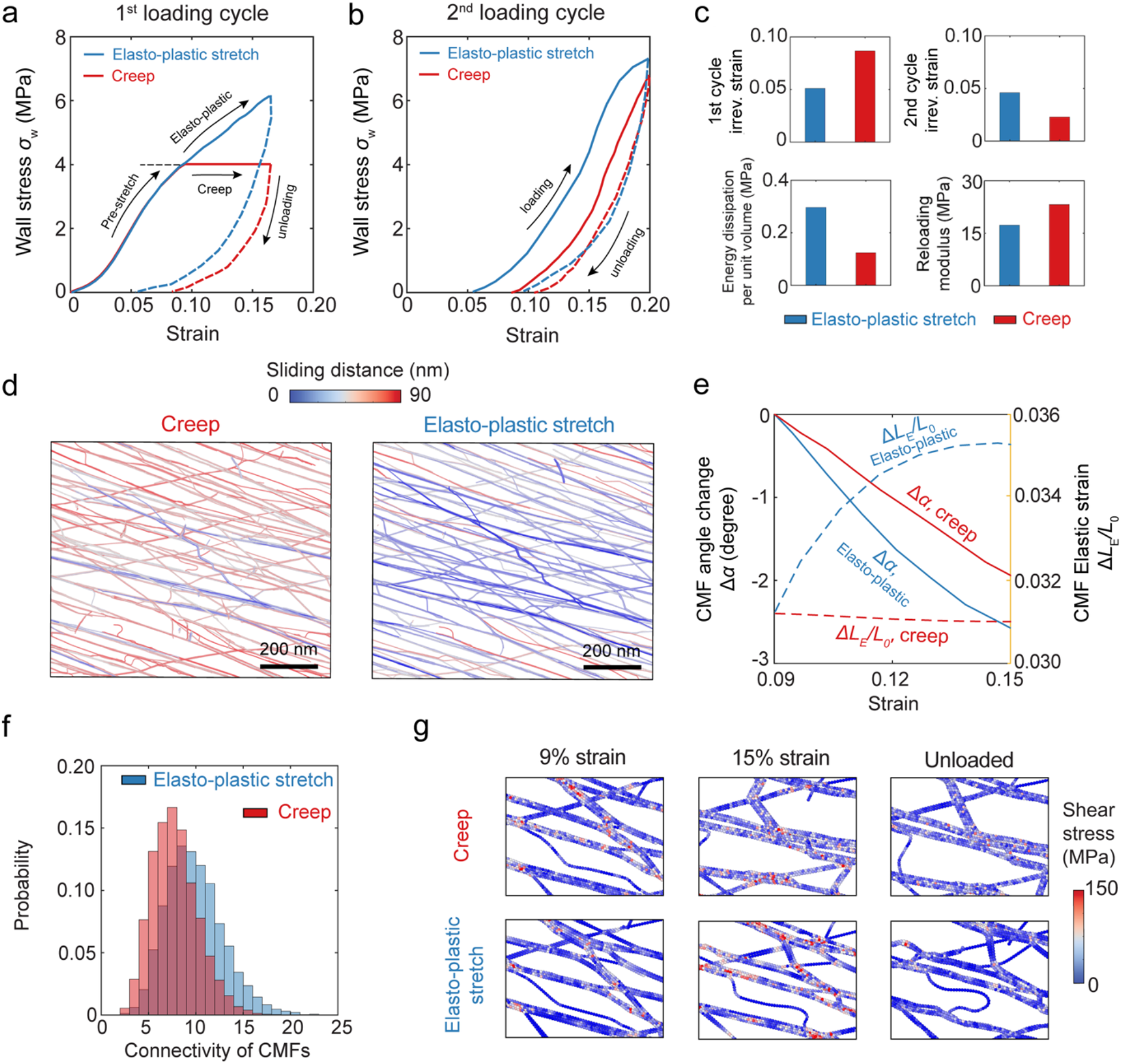
Cell walls after creep exhibit greater resistance to further plastic deformation. **(a-b)** Simulated stress-strain responses of cell walls after two cycles of loading (solid lines) and unloading (dashed lines). The stress-strain responses of the first (a) and the second (b) loading cycle are compared between elasto-plastic stretch (blue) and creep (red). In the first loading cycle, two identical walls are first pre-stretched to 4 MPa stress (~8% strain), followed by creep or elasto-plastic stretch to 15% strain. The walls are then elastically unloaded to zero stress. In the second loading cycle, the two walls are equally elasto-plastically stretched to 20% strain and unloaded. **(c)** Comparison of residual irreversible strain after the first and the second loading cycles, energy dissipation, and reloading modulus during the second loading cycle. **(d)** Heat map of irreversible sliding distances in paired CMFs during creep (left) and elasto-plastic stretch (right) simulations, at 15% tensile strain. The sliding distance is quantified as the summation of irreversible changes of registry position between interacting CMF pairs. Scale bar: 200 nm. **(e)** Change of the average CMF orientation angle Δα (solid lines) and the average elastic bond strain Δ*L*_E_/*L*_0_ (dashed lines) as a function of wall strain. **(f)** Comparison of CMF connectivity between creep and elasto-plastic stretch under 15% strain. **(g)** Representative microstructure changes during the first loading cycle of creep and elasto-plastic stretching. During creep, enhanced sliding promotes aggregation into thicker CMF bundles that persist after unloading and reloading, resulting in a more bundled microstructure than after rapid elasto-plastic stretching.

At the molecular level, the contrast in stress-strain responses noted above originates from different sliding patterns within CMF networks. At the initial stage of pre-stretch (0–3% wall strain), the CMF network deforms like a well-connected elastic lattice. At larger strains, collective sliding induces heterogeneous, time-dependent structural relaxation, leading to divergent stress-strain responses for slow creep versus rapid elasto-plastic stretch. Creep results in comparatively larger sliding distances at smaller stresses and disconnects parts of the CMF network, creating a visually more bundled and aligned network (**Fig. 4d**). CMF network structure after creep shows less elastic stretching and reorientation (**Fig. 4e**), consistent with published AFM measurements showing little CMF reorientation after creep^39^. Moreover, after creep the wall displayed lower connectivity (**Fig. 4f**) and more CMF bundling (**Fig. 4g**) compared with walls stretched elasto-plastically. These observations provide a microstructural basis for the reduced plastic deformation, lower hysteresis, and higher reloading stiffness observed after creep, as bundle reorganization suppresses further plastic deformation in subsequent loading cycles. The results highlight the potential structural advantage of slow creep of growing cell walls at moderate turgor pressure, compared with rapid plastic yielding.

## Discussion

Cell wall creep is central to plant growth, yet has long eluded a mechanically grounded molecular explanation and has often been conflated with plasticity, a second form of irreversible deformation. We show how wall creep may arise from stress-driven CMF sliding within the wall’s cellulose network, gated by discrete energy barriers stemming from noncovalent network formation of CMFs within the wall. Two sliding modes dominate: uniform sliding for short contacts and dislocation-mediated sliding for long contacts. The latter resembles the physics of polymer reptation^40^ and sets an upper bound on energy barriers for creep. Similar dislocation mechanisms may enable creep of other long-chain polymer networks stabilized by non-covalent interactions, such as polyethylene fibers^41^, wood^42^, fungal cell walls^43^, tendons^44^ and cuticles of molting crabs^45^. Other type of topological defects in cellulose networks such as twisting^46,47^, kinking^48^, or dislocations in smaller atomistic scales^49^, appear to follow similar nucleation-propagation cycles to facilitate irreversible deformation, yet their detailed defect dynamics needs future investigation.

Both creep and plasticity involve CMF sliding, the mode of which depends on contact length. In contrast to creep, plastic deformation involves the lowering of sliding energy barriers by wall stresses. Beyond a critical stress, the barriers collapse and CMFs undergo rapid sliding characteristic of plastic flow, which does not require thermal activation. Thus, creep and plastic deformation may share a common structural basis (CMF sliding) but differ sharply in kinetics, stress thresholds, and rearrangements of microfibril networks—features absent from growth models that treat wall enlargement as purely elasto-plastic deformation. This contrast also highlights a potential advantage of creep over plasticity: it enables sustained enlargement while preserving structural integrity of the wall.

We note that the current model simplifies wall structure to its three major components, neglects cellulose polymorph-specific surface energetics and does not include the action of expansins. Nevertheless, this framework suggests testable hypotheses for α-expansin-induced wall creep^18,50,51^, a long-standing question in plant biology. Expansins may target energetic “hot spots”^18^ within the cellulose network such as CMF ends or junctions or noncrystalline regions, where they might nucleate dislocation-like defects that promote sliding far from the nucleation site. Such a scheme could account for the effectiveness of α-expansins at low concentrations in growing walls^18^. While dislocations and their gliding kinetics are well established as the basis of plasticity and creep in inorganic crystals^52–55^, dislocation dynamics in semi-crystalline polymers^56^, including cellulose networks, have so far received remarkably little attention. Recognizing dislocation-mediated motion in plant cell walls reframes wall biomechanics and highlights an overlooked role for polymeric dislocations in semicrystalline biological materials. These insights suggest new avenues for probing expansin function, experimentally testing dislocation dynamics, and applying defect engineering concepts^57^ to tune the mechanics and growth behavior of cellulose-based and other fibrous networks.

## Methods

### Creep tests

Cell wall specimens consisted of a 7-μm-thick strip of outer epidermal wall isolated from onion bulbs (*Allium cepa*) as described previously^14^. Strips of outer periclinal cell walls (10 mm × 3 mm × 7 μm) were peeled from the center region of the abaxial side of the fifth fleshy scale (counting from the outermost layer). Wall samples were incubated with 5 mg mL^-1^ Pronase in 20 mM HEPES (pH 7.0) with agitation at 26°C for 1.5 h to remove endogenous expansin activity. With custom creep devices^34^, wall strips were initially clamped (5 mm between clamps) under a pre-load of 0.6 g (in 20 mM HEPES pH 7.0 buffer) to enable accurate initial position measurements. Two minutes after clamping, the wall strips were further loaded to desired levels and changes in length were recorded every 30 s for 1.5 h. Wall creep was assessed at loads of 2, 4, 6, 8, 10, 12, 14, and 16 grams with two wall samples from three onions (n = 6).

### Stress and shear force calculations

We computed the bead-level stress tensor *σ*_*ij*_ by the virial stress formulation. The stress tensor can be decomposed into its spherical component −*p***I** and deviatoric component *τ*_*ij*_ (shear stress), given by 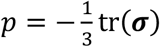 and *τ*_*ij*_ = *σ*_*ij*_ + 3*p*δ_*kk*_, where δ_*ij*_ is the Kronecker delta. The driving force of sliding can be considered as the shear force parallel to the interfaces of CMFs. The magnitude of the interfacial shear force ***f***^(*k*)^ on bead *k* can be calculated by

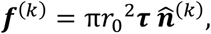

where 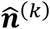 is the normal direction of the cellulose-cellulose interface. The average interfacial shear stress on the *i*-th CMF can then be defined as

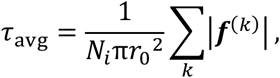

where *N*_*i*_ is the total number of beads in the *i*-th CMF chain, and *r*_0_ is the radius of coarse-grained beads.

### NEB calculations

To estimate the free energy barrier and to find the minimum energy path for CMF sliding, we adopted a two-step NEB calculation to separate the dislocation nucleation (the first step) and dislocation gliding processes (the second step). During each step, two configurations of CMF pairs were generated and relaxed using sufficiently long dynamic equilibration (10 ns), used as the initial and final configurations of minimum energy paths respectively. During the dynamic equilibration, one end bead was fixed and a constant force *F* = *σ*_f_π*R*^2^ is applied on another end bead, parallel to the direction of CMF pairs. Then, the two fully relaxed geometries are taken as the endpoints of NEB simulations. For the dislocation nucleation process, the initial configuration had contact length *L* = 3*n* nm, where *n* is the number of overlapping beads. The final configuration was obtained by compressing one free end with one-bead distance to its nearest registry, then select an energetically optimal wavelength. For the dislocation gliding process, the initial configuration was the fully nucleated dislocation, and the final configuration was obtained by moving the center of dislocation by one registry. The intermediate images were initialized by linearly interpolating the bead positions between two endpoints, using 30 images in total. The quick-min optimizer^58^ was used to relax the initial reaction pathway to the minimum energy path. NEB calculations were considered converged if the force component perpendicular to the elastic band was below 3 × k0^-4^ pN. The potential energy associated with the applied constant force *F* was included during the minimization. The parallel and perpendicular spring nudging constant were *K*_∥_ = 20 *k*_=_*T*/Å^2^ and *K*_⊥_ = 2 *k*_=_*T*/Å^2^, respectively. Both the initial equilibration and the NEB calculations were performed under a two-dimensional constraint.

### Definition of sliding coordinates

Sliding coordinates were introduced to quantify the relative position of transition states along the minimum-energy path (MEP) using a single scalar variable. They were defined from the cumulative geodesic distances along the MEP obtained by NEB simulations. For an intermediate state *i*, the incremental path length along MEP is 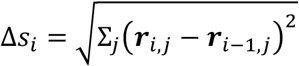, where ***r*** is the position of bead *j* in state *i*. The reaction coordinate *s*_*i*_ is then obtained as the cumulative sum along the MEP, 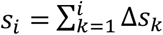, and normalized such that *s* = *s*_0_ corresponds to the initial state and *s* = *s*_0_ + k to the final state. For uniform sliding, *s*_0_ = 0. For dislocation-mediated sliding, *s*_0_ = 0 indicates the beginning of dislocation nucleation, and *s*_0_ = k indicates the beginning of dislocation gliding.

### CGMD-MC algorithm

Monte Carlo moves were defined as unidirectional sliding events on each CMF pair *i* and *j*, with sliding strain rates governed by the Arrhenius-type kinetics 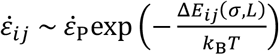, where 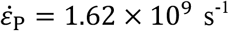 was fitted from the experimental creep response. Note that following de Gennes^40^ we neglect reverse sliding because CMFs buckle under compression. The energy barrier Δ*E*_*ij*_(*σ, L*) was obtained from NEB simulations (**Fig. 2e**). We applied a rejection-free kinetic Monte Carlo algorithm^59^ to all CMF pairs and defined the local sliding direction 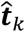 on each bead as 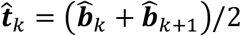, where 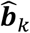 and 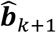 are the directions of the two bonds connected to the bead *k*, respectively. The sign of 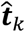 was determined by 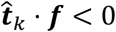, which means the sliding direction should oppose the interfacial shear force. During a single sliding event, the position of beads ***r***_k_ was updated by 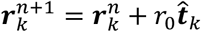, where r0 is the equilibrium bond length, and the superscript *n* denotes the *n*-th timestep. We further assumed sliding events only extend but not shrink CMF overlap, and this unidirectional assumption was enforced by calculating the overall direction of shear forces. Under these assumptions, the transition rate of a single sliding event *i* is given by 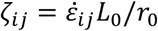.

The coupled CGMD-MC algorithm consisted of an inner CGMD loop and an outer kinetic Monte Carlo loop. The inner CGMD loop handled transient dynamics to keep the system near local equilibrium, whereas the outer loop applied the sliding kinetics to extend the simulation to much longer timescales. In the outer loop, all possible sliding pairs were identified and their sliding rates are calculated by *ζ*_*ij*_. The cumulative updating rate was then given by 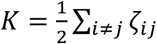 A small subset of CMF pairs was randomly sampled by ranking *ζ*_*ij*_ (**Supplementary Note 2**). Then, the corresponding sliding event was then applied to the chosen CMF pair, and the simulation clock was advanced by Δ*t* = *MK*^-1^, where *M* is the number of picked CMF pairs during each sampling. After the sliding event, the inner loop ran 500 steps of molecular dynamics time integration to re-equilibrate the system. We implemented the CGMD-MC algorithm based on the open-source molecular dynamics package LAMMPS. A custom fix style command was implemented to insert MC calculations during molecular dynamics time integration.

### Definition of CMF connectivity

CMF connectivity was defined as the number of noncovalent contacts *N*_c,*I*_ on the *i*-th microfibril. The average CMF connectivity was then defined as 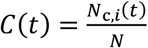, where *N* is the total number of CMFs. During CGMD-MC simulations, an adjacency map was maintained to record the neighbor information in CMF networks, and *N*_c,*i*_(*t*) was calculated by traversing the adjacency map at each timestep.

## Data and code availability

The code used for CGMD-MC simulations and NEB simulations is publicly available on Zenodo (10.5281/zenodo.19208520). Source data are provided with this paper.

## Acknowledgments

We thank Liza Wilson for providing the AFM image of onion cell walls, as well as Luyi Feng and Prof. Adri van Duin for discussing the nudged elastic band simulations. We thank Prof. Yao Zhang for discussing the assembly of the initial cell wall configurations. We thank Prof. Enrico Coen, Prof. Charlie Anderson, and Prof. Spencer Szczesny for discussions of the results and the manuscript. Research reported in this publication was supported by the Center for Engineering MechanoBiology (CEMB), an NSF Science and Technology Center, under grant agreement CMMI: 15-48571. We acknowledge the support of computational resources from the Advanced Cyberinfrastructure Coordination Ecosystem: Services & Support (ACCESS) program, which is supported by U.S. National Science Foundation grants #2138259, #2138286, #2138307, #2137603, and #2138296.

## Author contributions

Conceptualization: D.J.C., J.H. and S.Z.

Formal Analysis: C.L., J.Y., D.J.C. and S.Z.

Funding Acquisition: D.J.C. and S.Z.

Investigation: C.L., J.Y., D.J.C. and S.Z.

Methodology: C.L., J.Y., D.J.C. and S.Z.

Software: C.L. and S.Z.

Supervision: D.J.C. and S.Z.

Visualization: C.L.

Writing – original draft: C.L., D.J.C. and S.Z.

Writing – review & editing: C.L., J.Y., J.H., D.J.C. and S.Z.

## Competing interests

The authors declare that they have no competing interests.

## Supplementary information

Supplementary information is available for this paper online.

Correspondence and requests for materials should be addressed to either.

## Notes

### Competing Interest Statement

The authors have declared no competing interest.

## References

1 Cosgrove, D. J. Structure and growth of plant cell walls. Nature Reviews Molecular Cell Biology 25, 340–358 (2024).

2 Braidwood, L., Breuer, C. & Sugimoto, K. My body is a cage: mechanisms and modulation of plant cell growth. New Phytologist 201, 388–402 (2014).

3 Coen, E. & Cosgrove, D. J. The mechanics of plant morphogenesis. Science 379, eade8055 (2023).

4 Delmer, D., Dixon, R. A., Keegstra, K. & Mohnen, D. The plant cell wall—dynamic, strong, and adaptable—is a natural shapeshifter. The Plant Cell 36, 1257–1311 (2024).

5 Fratzl, P. & Weinkamer, R. Nature’s hierarchical materials. Progress in Materials Science 52, 1263–1334 (2007).

6 Zhang, J. et al. Toughness enhancement by massive dislocation absorption at the crack front. Proceedings of the National Academy of Sciences 122, e2511830122 (2025).

7 Ashby, M. F. A first report on deformation-mechanism maps. Acta Metallurgica 20, 887–897 (1972).

8 Zhang, Y. et al. Molecular insights into the complex mechanics of plant epidermal cell walls. Science 372, 706–711 (2021).

9 Parre, E. & Geitmann, A. Pectin and the role of the physical properties of the cell wall in pollen tube growth of Solanum chacoense. Planta 220, 582–592 (2005).

10 Chen, S. et al. Fibrous network nature of plant cell walls enables tunable mechanics for development. Nature Communications 16, 7565 (2025).

11 Geitmann, A., Mulder, B. M., Persson, S. & Spalding, E. P. Modeling the molecular structures and dynamics responsible for the remarkable mechanical properties of a plant cell wall. Faculty Review 11, 24 (2022).

12 Silveira, S. R. et al. Mechanical interactions between tissue layers underlie plant morphogenesis. Nature Plants 11, 909–923 (2025).

13 Burton, R. A., Gidley, M. J. & Fincher, G. B. Heterogeneity in the chemistry, structure and function of plant cell walls. Nature Chemical Biology 6, 724–732 (2010).

14 Yu, J., Zhang, Y. & Cosgrove, D. J. The nonlinear mechanics of highly extensible plant epidermal cell walls. Proceedings of the National Academy of Sciences 121, e2316396121 (2024).

15 Mosca, G. et al. Growth and tension in explosive fruit. Current Biology 34, 1010-1022. e1014 (2024).

16 Zhang, T., Tang, H., Vavylonis, D. & Cosgrove, D. J. Disentangling loosening from softening: insights into primary cell wall structure. The Plant Journal 100, 1101–1117 (2019).

17 Zhang, Y., Yu, J. & Cosgrove, D. J. Mechanical Roles of Polysaccharide Assembly and Interactions in Plant Cell Walls. Biomacromolecules (2025).

18 Cosgrove, D. J. Plant cell wall loosening by expansins. Annual Review of Cell and Developmental Biology 40 (2024).

19 Samalova, M., Gahurova, E. & Hejatko, J. Expansin-mediated developmental and adaptive responses: A matter of cell wall biomechanics? Quantitative Plant Biology 3, e11 (2022).

20 Zeng, H. Y. et al. Origin and evolution of auxin-mediated acid growth. Proceedings of the National Academy of Sciences 121, e2412493121 (2024).

21 Dror, R. O., Dirks, R. M., Grossman, J., Xu, H. & Shaw, D. E. Biomolecular simulation: a computational microscope for molecular biology. Annual Review of Biophysics 41, 429–452 (2012).

22 Zuckerman, D. M. Equilibrium sampling in biomolecular simulations. Annual Review of Biophysics 40, 41–62 (2011).

23 Plimpton, S. Computational limits of classical molecular dynamics simulations. Computational Materials Science 4, 361–364 (1995).

24 Zhang, T., Vavylonis, D., Durachko, D. M. & Cosgrove, D. J. Nanoscale movements of cellulose microfibrils in primary cell walls. Nature Plants 3, 1–6 (2017).

25 Eyring, H. Viscosity, Plasticity, and Diffusion as Examples of Absolute Reaction Rates. The Journal of chemical physics 4, 283–291 (1936).

26 Jiang, Y. et al. Activation Volume Facilitating Chemical Reaction under Mechanical Stress. Physical Review Letters 135, 128001 (2025).

27 Jónsson, H., Mills, G. & Jacobsen, K. W. in Classical and Quantum Dynamics in Condensed Phase Simulations 385–404 (World Scientific, 1998).

28 Cox, H. L. The elasticity and strength of paper and other fibrous materials. British Journal of Applied Physics 3, 72 (1952).

29 Keckes, J. et al. Cell-wall recovery after irreversible deformation of wood. Nature Materials 2, 810–814 (2003).

30 Mittal, N. et al. Multiscale Control of Nanocellulose Assembly: Transferring Remarkable Nanoscale Fibril Mechanics to Macroscale Fibers. ACS Nano 12, 6378–6388 (2018).

31 Kretschmann, D. Natural materials - Velcro mechanics in wood. Nature Materials 2, 775–776 (2003).

32 Schulze, T. P. Efficient kinetic monte carlo simulation. Journal of Computational Physics 227, 2455–2462 (2008).

33 Bernardi, R. C., Melo, M. C. & Schulten, K. Enhanced sampling techniques in molecular dynamics simulations of biological systems. Biochimica et Biophysica Acta (BBA)-General Subjects 1850, 872–877 (2015).

34 Durachko, D. M., Park, Y. B., Zhang, T. & Cosgrove, D. J. Biomechanical characterization of onion epidermal cell walls. Bio-protocol 7, e2662–e2662 (2017).

35 Green, P., Erickson, R. & Buggy, J. Metabolic and physical control of cell elongation rate: in vivo studies in Nitella. Plant Physiology 47, 423–430 (1971).

36 Lockhart, J. A. An analysis of irreversible plant cell elongation. Journal of Theoretical Biology 8, 264–275 (1965).

37 Chakraborty, J., Luo, J. & Dyson, R. J. Lockhart with a twist: Modelling cellulose microfibril deposition and reorientation reveals twisting plant cell growth mechanisms. Journal of Theoretical Biology 525, 110736 (2021).

38 Duffo, P., Monasse, B., Haudin, J. M., G’Sell, C. & Dahoun, A. Rheology of polypropylene in the solid state. Journal of Materials Science 30, 701–711 (1995).

39 Marga, F., Grandbois, M., Cosgrove, D. J. & Baskin, T. I. Cell wall extension results in the coordinate separation of parallel microfibrils: evidence from scanning electron microscopy and atomic force microscopy. The Plant Journal 43, 181–190 (2005).

40 De Gennes, P.-G. Reptation of a polymer chain in the presence of fixed obstacles. The Journal of Chemical Physics 55, 572–579 (1971).

41 O’Connor, T. C. & Robbins, M. O. Molecular models for creep in oriented polyethylene fibers. The Journal of Chemical Physics 153, 144904 (2020).

42 Stevanic, J. S. & Salmen, L. Molecular origin of mechano-sorptive creep in cellulosic fibres. Carbohydrate Polymers 230 (2020).

43 Chakraborty, A. et al. A molecular vision of fungal cell wall organization by functional genomics and solid-state NMR. Nature Communications 12, 6346 (2021).

44 Wang, X. T. & Ker, R. F. Creep rupture of wallaby tail tendons. Journal of Experimental Biology 198, 831–845 (1995).

45 Chen, P.-Y., Lin, A. Y.-M., McKittrick, J. & Meyers, M. A. Structure and mechanical properties of crab exoskeletons. Acta Biomaterialia 4, 587–596 (2008).

46 Usov, I. et al. Understanding nanocellulose chirality and structure–properties relationship at the single fibril level. Nature Communications 6, 7564 (2015).

47 Savin, A., Khalack, J., Christiansen, P. L. & Zolotaryuk, A. V. Twisted topological solitons and dislocations in a polymer crystal. Physical Review B 65, 054106 (2002).

48 Ciesielski, P. N. et al. Nanomechanics of cellulose deformation reveal molecular defects that facilitate natural deconstruction. Proceedings of the National Academy of Sciences 116, 9825–9830 (2019).

49 Lipchinsky, A. How do expansins control plant growth? A model for cell wall loosening via defect migration in cellulose microfibrils. Acta Physiologiae Plantarum 35, 3277–3284 (2013).

50 McQueen-Mason, S., Durachko, D. M. & Cosgrove, D. J. Two endogenous proteins that induce cell wall extension in plants. The Plant Cell 4, 1425–1433 (1992).

51 McQueen-Mason, S. J. & Cosgrove, D. J. Expansin mode of action on cell walls (analysis of wall hydrolysis, stress relaxation, and binding). Plant Physiology 107, 87–100 (1995).

52 Weertman, J. Theory of steady-state creep based on dislocation climb. Journal of Applied Physics 26, 1213–1217 (1955).

53 Peach, M. & Koehler, J. The forces exerted on dislocations and the stress fields produced by them. Physical Review 80, 436 (1950).

54 Dong, L. et al. Borrowed dislocations for ductility in ceramics. Science 385, 422–427 (2024).

55 Masters, B. Dislocation loops in irradiated iron. Nature 200, 254–254 (1963).

56 Bartczak, Z., Galeski, A., Argon, A. S. & Cohen, R. E. On the plastic deformation of the amorphous component in semicrystalline polymers. Polymer 37, 2113–2123 (1996).

57 Jangizehi, A., Schmid, F., Besenius, P., Kremer, K. & Seiffert, S. Defects and defect engineering in Soft Matter. Soft Matter 16, 10809–10859 (2020).

58 Sheppard, D., Terrell, R. & Henkelman, G. Optimization methods for finding minimum energy paths. The Journal of Chemical Physics 128 (2008).

59 Hatch, H. W., Corti, D. S., Kofke, D. A. & Shen, V. K. Best Practices for Developing Monte Carlo Methodologies in Molecular Simulations [Article v1. 0]. Living Journal of Computational Molecular Science 6, 3289–3289 (2025).

