## Supplementary Text for "Collective microfibril sliding underlies plant cell wall creep"

**This PDF file includes:**

Extended Data Figures 1 to 6
Supplementary Notes 1 to 2
Table S1
Video captions S1 to S4
References

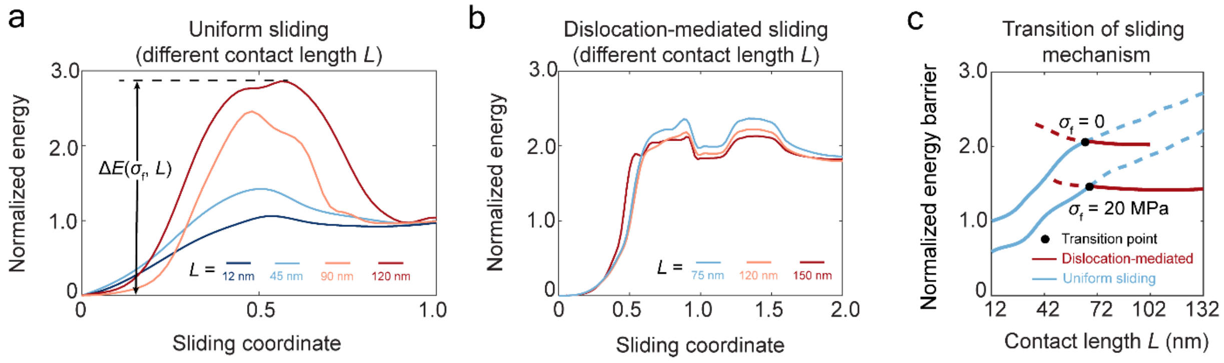


**Extended Data Fig. 1 | Different energetic scaling leading to the transition of sliding mechanisms. (a)** Minimum energy paths of uniform sliding under different contact lengths. The sliding energy barrier monotonically increases with contact length, because CMF pairs are completely detached during uniform sliding. **(b)** Minimum energy paths of dislocation-mediated sliding under different contact length. The energy barrier keeps nearly unchanged under long contact lengths, because dislocation nucleation events are localized and independent of contact length. Energies are normalized by $2D_{01} = 90\ k_{\mathrm{B}}T$, where $D_{01}$ is the non-bonded interaction strength between two beads. **(c)** Normalized sliding energy barriers of uniform sliding (blue lines) and dislocation-mediated sliding (red lines), under different contact lengths and stresses. Solid lines denote paths with minimal energy barriers, and dashed lines denote alternative paths with higher energy barriers. Minimal energy paths can be determined from the intersections (black dots) between blue and red lines. The intersections denote the transition from uniform sliding to dislocation-mediated sliding, arising from different scaling of energy barriers.

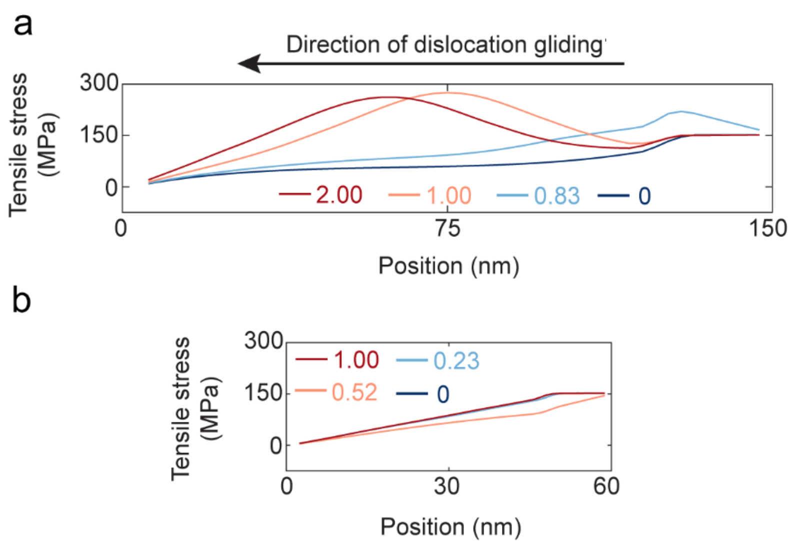


**Extended Data Fig. 2 | Distinct evolution of CMF tensile stresses for different sliding mechanisms. (a)** Tensile stresses for dislocation-mediated sliding ($L$ = 150 nm, $\sigma_f$ = 150 MPa) at different sliding coordinates. A dislocation first nucleates at the right end (sliding coordinate from 0 to 1.0), leading to a steeper gradient of tensile stresses. Then the dislocation starts to glide into the chain (sliding coordinate from 1.0 to 2.0), visualized by the moving earthworm-like profiles of tensile stresses. During dislocation gliding, the tensile (and shear) stresses are not symmetric around the dislocation center, driving unidirectional gliding. **(b)** Tensile stresses for uniform sliding ($L$ = 60 nm, $\sigma_f$ = 150 MPa) at different sliding coordinates. The tensile stress distribution shows minimal changes during sliding, suggesting that the deformation mode of the chain remains unchanged, consistent with the concept of uniform sliding.

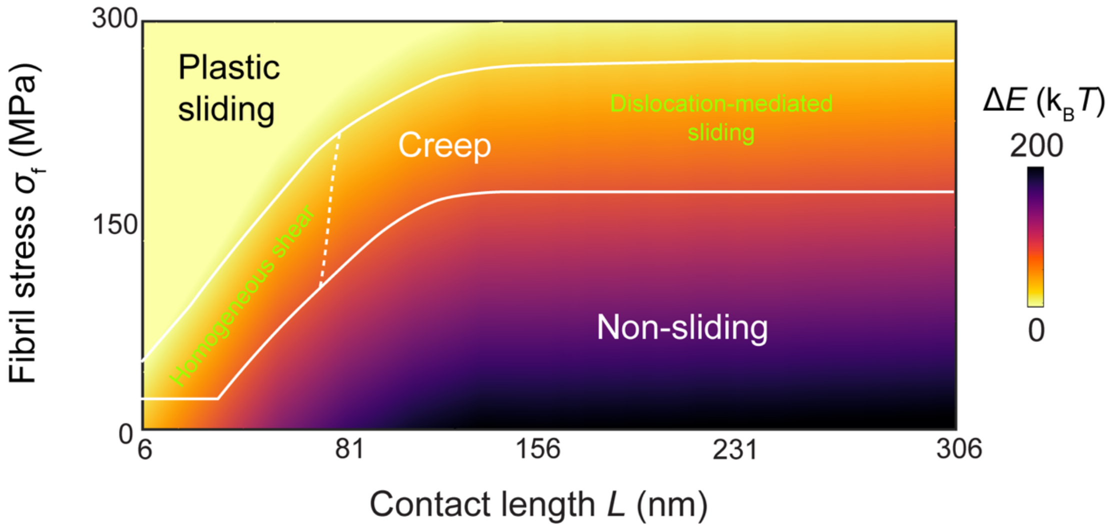


**Extended Data Fig. 3 | Extended phase diagram of sliding free energy barriers.** Energies are normalized by $2D_{01} = 90\ k_{\mathrm{B}}T$. Regions of non-sliding (elastic deformation only), creep, and plasticity are separated by white solid lines. The creep region is further separated by two distinct sliding mechanisms: uniform sliding at low contact lengths and dislocation-mediated gliding at high contact lengths.

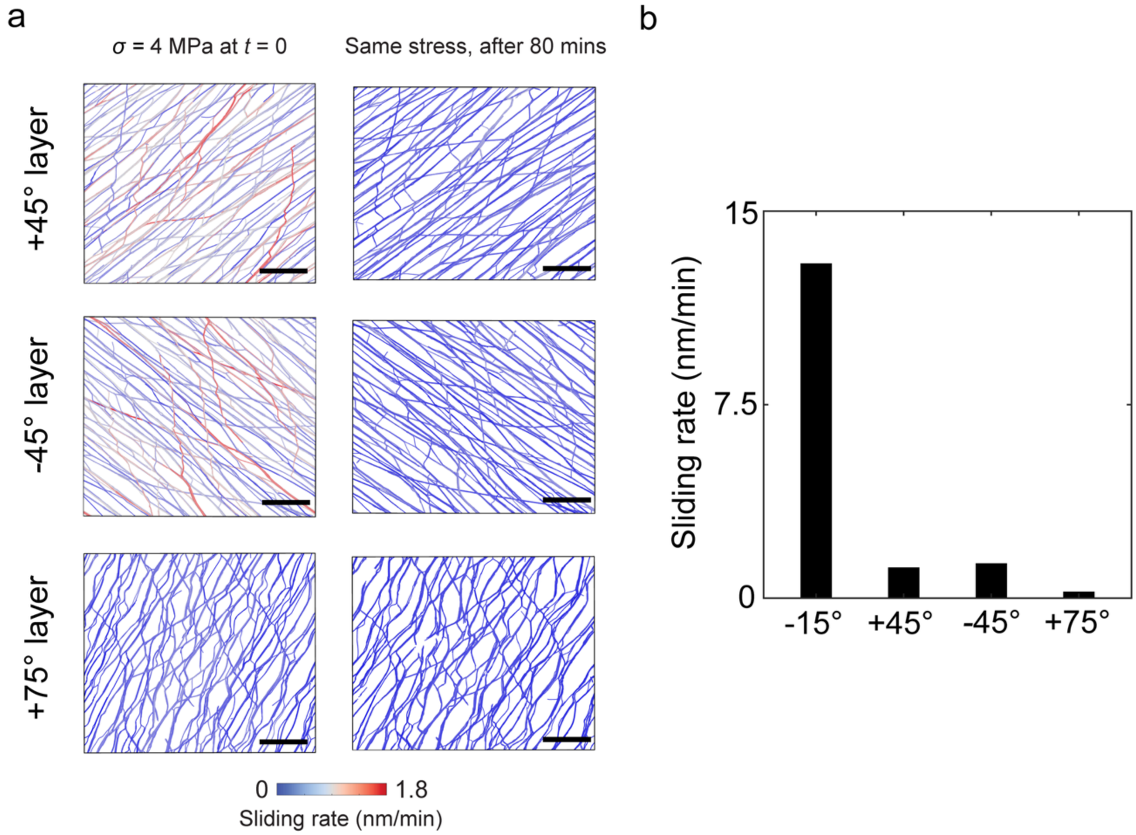


**Extended Data Fig. 4 | Lamellae not parallel to the stretching direction exhibit much less CMF sliding.** **(a)** Structural changes of −45°, +45°, and +75° lamellae during creep, colored by CMF sliding rates. Scale bar: 200 nm. **(b)** Average sliding rate over all CMFs in each layer, at $t = 0$.

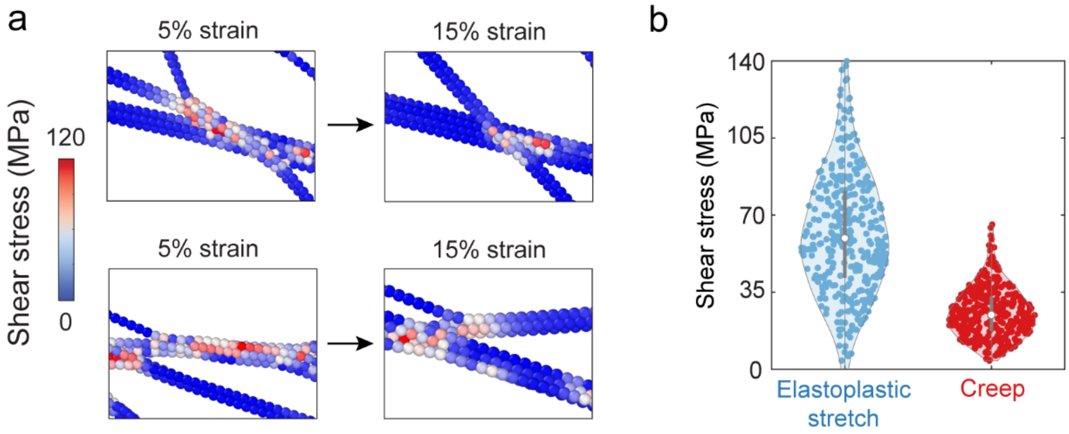


**Extended Data Fig. 5 | Creeping walls show frequent changes of bundling that lead to stress redistribution. (a)** Two examples of bundling changes during creep, color-coded by the shear stress magnitude. Both top and bottom examples show bundle merging and splitting during collective sliding, along with CMF stretching and reorientation. **(b)** Distributions of chain-averaged shear stress under 15% strain, where each scatter represents a single CMF in the −15° lamella.

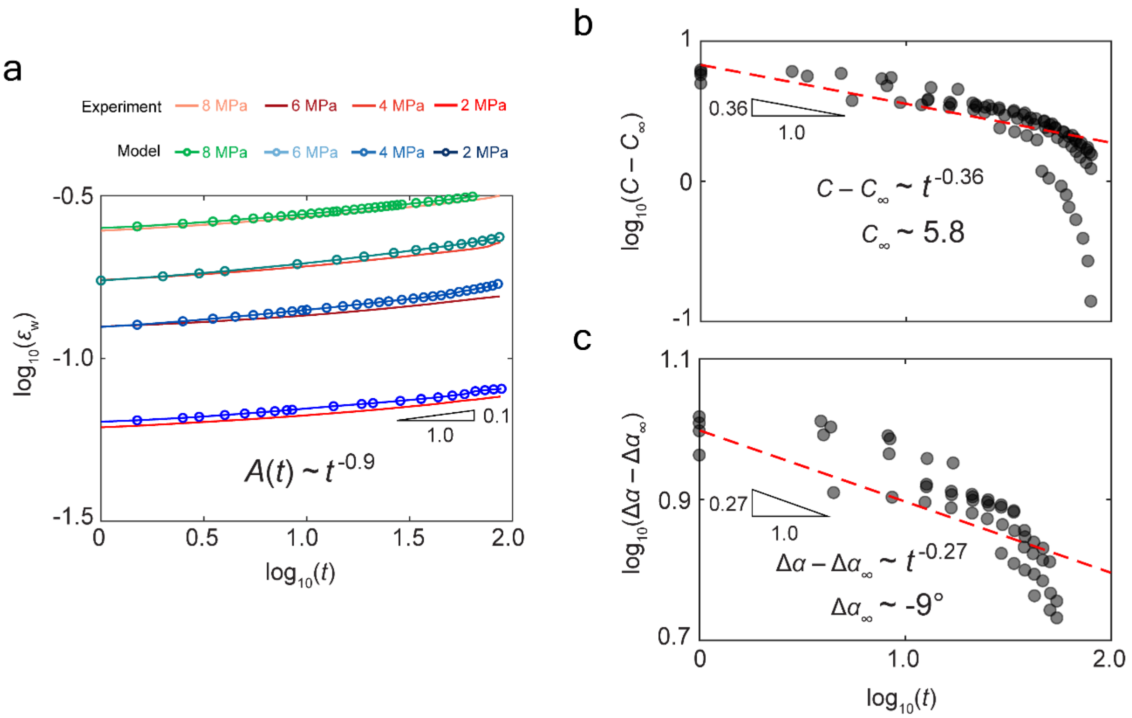


**Extended Data Fig. 6 | Empirical creep laws fitted from stresses and structural changes of CMF networks. (a)** Experimentally measured (solid lines) and simulated (connected scatters) time evolution of wall strain under different stresses ($\sigma_w$ = 2, 4, 6, 8 MPa) on a log-log plot. The stress exponent can be fitted by assuming $\dot{\varepsilon}_w \sim A(t)(\sigma_w - Y)^n = t^m(\sigma_w - Y)^n$, where $m$ and $n$ are time and stress exponents respectively. After taking the logarithm on both sides, linear regression gives $m \sim -0.9$ and $n \sim 1.1$ ($R^2$ = 0.961 and 0.976 respectively), leading to a power law $A(t) \sim t^{-0.9}$ that quantitatively captures the evolution of structure-related factor under different stresses. **(b-c)** Simulated time evolution (black scatters) of connectivity $C(t)$ in (b) and CMF angle change $\Delta\alpha(t)$ in (c). Red dashed lines denote the power-law fitted results. Altogether, assuming $A(t) \sim (C - C_\infty)^p(\Delta\alpha - \Delta\alpha_\infty)^q(\sigma_w - Y)^n$, nonlinear regression gives $p$ = 2.1, $q \sim$ 0.4, $n \sim$ 1.1, leading to a creep law $\dot{\varepsilon}_w \sim (C - 5.8)^{2.1}(\Delta\alpha + 9°)^{0.4}(\sigma_w - 0.25\ \text{MPa})^{1.1}$ which does not explicitly contain time.

**Supplementary Notes**

### *1. Coarse-grained molecular dynamics (CGMD) model of plant cell wall*

Zhang et al.[1] developed a coarse-grained molecular dynamics (CGMD) model to recapitulate the mesoscale structure and complex mechanical properties of plant cell walls, including nonlinear elasticity, plasticity and hysteresis. The model reveals different microstructural deformation modes contributing to tensile mechanics of cell walls. Details of this CGMD model are elaborated in the Supplementary Information of the reference[1]. We adopt the initial configuration and model parameters without modification.

### *2. Monte Carlo (MC) sampling algorithm*

As we emphasized in the main text, the sliding process between two contacting CMF chains is a sequence of alternating dislocation nucleation and gliding events. The sliding energy barrier is dominated by the barrier of nucleation, which is a function of applied stresses and the contact length. Given the sliding barrier $\Delta E(\sigma, L)$ of a pair of CMFs, one can apply transition state theory[2] (TST) $\dot{\varepsilon} \sim \dot{\varepsilon}_{\mathrm{P}} \exp\left(-\frac{\Delta E(\sigma,L)}{k_{\mathrm{B}}T}\right)$ to calculate the sliding rate of all CMF pairs in the cell wall model. Here we describe the kMC time stepping algorithm as follows:

*Step 1*: At time $t$, read the cell wall configuration $\{(\boldsymbol{x}_k, \boldsymbol{v}_k)\}$, where $\boldsymbol{x}_k$ and $\boldsymbol{v}_k$ are locations and velocities for the $k$-th bead. For each CMF pair $i$ and $j$, construct an adjacency matrix $A_{ij} = 1$ or 0 denoting whether the CMF $i$ and $j$ are interacting.

*Step 2*: For those interacting CMF pairs, calculate the sliding energy barrier $\Delta E_{ij}(\sigma, L)$ by interpolating the NEB results. The sliding rate is given by $\dot{\varepsilon}_{ij} = \dot{\varepsilon}_{\mathrm{P}} \exp\left(-\frac{\Delta E_{ij}(\sigma,L)}{k_{\mathrm{B}}T}\right)$, where $T = 300$ K.

*Step 3*: For each CMF pair $i$ and $j$, calculate the transition rate between the states as $\zeta_{ij} = \dot{\gamma}_{ij} L / r_0$. Then, sort $\zeta_{ij}$ and store them in a one-dimensional array. Calculate the cumulative transition rate $K = \frac{1}{2}\Sigma_{i \neq j} \zeta_{ij}$.

*Step 4*: Randomly select $M$ CMF pairs, such that each random CMF pair $\zeta_{ij}$ is sampled by a uniformly distributed random number $z \in (0,1]$, by the criteria that $\zeta_{ij} < zK < \zeta_{pq}$.

*Step 5*: For the selected $M$ CMF pairs, the timestep can be given by $\Delta t = MK^{-1}$. Update the time to $t + \Delta t$. Update the position of the selected CMF pair by the rule $\boldsymbol{r}_i^{n+1} = \boldsymbol{r}_i^n + r_0 \hat{\boldsymbol{t}}_i$, where $r_0$ is the equilibrium bond length, and the superscript $n$ denotes the $n$-th kMC timestep. The local sliding direction

$\hat{\boldsymbol{t}}_i$ on each bead is given by $\hat{\boldsymbol{t}}_i = (\hat{\boldsymbol{b}}_i + \hat{\boldsymbol{b}}_{i+1})/2$, where $\hat{\boldsymbol{b}}_i$ and $\hat{\boldsymbol{b}}_{i+1}$ are the directions of the two bonds connected to the bead $i$, respectively.

*Step 6*: Perform $N_{\mathrm{MD}}$ steps of molecular dynamics (NPT ensemble for creep simulations, NVT ensemble for stress relaxation simulations) to relax the system to a new mechanical equilibrium at given boundary conditions (constant force for creep simulations, constant displacement for stress relaxation simulations). Return to *Step 1* until the time upper limit is reached.

The above rejection-free Monte Carlo algorithm, the rate constant of which is approximated by harmonic transition state theory[3], was first proposed by the reference[4]. Importantly, one can show that the probability that a unit process in a continuous-time Markov process has not occurred by time $t$ is $p_{\mathrm{survival}}(t) = \exp(-\zeta t)$. The probability distribution for the first passage time is then given by:

$$p(t) = \frac{\mathrm{d}}{\mathrm{d}t}\left(1 - p_{\mathrm{survival}}(t)\right) = \zeta \exp(-\zeta t), \tag{S12}$$

and the average first passage time $t_0$ is given by

$$t_0 = \int_0^{+\infty} t\, p(t)\, \mathrm{d}t = \frac{1}{\zeta}, \tag{S13}$$

where the rate $\zeta$ can further be approximated by the Arrhenius law $\zeta \sim \exp\left(\frac{-\Delta E}{k_{\mathrm{B}}T}\right)$. It then follows that the characteristic timescale for a unit process, simulated by the above MC algorithm, is $t_0 \sim \exp\left(\frac{\Delta E}{k_{\mathrm{B}}T}\right)$. The MC parameters are listed in **Table S1**.

**Table S1. Parameters used in Monte Carlo simulations.**

| $\dot{\varepsilon}_{\mathrm{P}}$ | $T$ | $M$ | $N_{\mathrm{MD}}$ |
|---|---|---|---|
| $1.62 \times 10^9$ s$^{-1}$ | 300 K | 5 | 500 |

**Video captions**

**Video S1**: **Homogeneous sliding.** Under short contact length ($L$ = 60 nm), homogeneous sliding crosses a single energy barrier, dominated by the energy needed to disconnect the entire contact surface. The video shows all NEB intermediate stages (colored by interfacial shear stresses) and the corresponding energy landscape. Scale bar: 18 nm.

**Video S2**: **Dislocation-mediated sliding.** Under long contact length ($L$ = 150 nm), dislocation-mediated sliding first crosses a major dislocation nucleation energy barrier, then multiple minor dislocation gliding barriers. The video first shows NEB intermediate stages (colored by interfacial shear stresses) and the energy landscape for dislocation nucleation, then shows the consequent two dislocation gliding events. Note that each single gliding process is denoted by advancing the sliding coordinate by 1.0. CMF chains are colored by interfacial shear stress. Scale bar: 18 nm.

**Video S3**: **Cell wall creep.** The slowly creeping wall exhibits enhanced CMF sliding, leading to more bundling from disconnected CMF contacts. The video shows simulated structural changes of CMF networks (900 nm × 900 nm × 160 nm) during creep (0–80 min, pre-stretched by ~4 MPa external stress), colored by the magnitude of interfacial shear stresses. Scale bar: 200 nm.

**Video S4: Cell wall elasto-plastic stretch.** Rapid elasto-plastically stretched wall shows increased CMF shear stresses, more CMF reorientation, and less disconnected CMF contacts. The video shows simulated structural changes of CMF networks (900 nm × 900 nm × 160 nm) during elastoplastic stretch (0–15% strain), colored by the magnitude of interfacial shear stresses. Scale bar: 200 nm.

**References**


1 Zhang, Y. *et al.* Molecular insights into the complex mechanics of plant epidermal cell walls. *Science* **372**, 706-711 (2021).

2 Eyring, H. Viscosity, plasticity, and diffusion as examples of absolute reaction rates. *The Journal of Chemical Physics* **4**, 283-291 (1936).

3 Truhlar, D. G., Garrett, B. C. & Klippenstein, S. J. Current status of transition-state theory. *The Journal of Physical Chemistry* **100**, 12771-12800 (1996).

4 Bortz, A. B., Kalos, M. H. & Lebowitz, J. L. A new algorithm for Monte Carlo simulation of Ising spin systems. *Journal of Computational Physics* **17**, 10-18 (1975).
